# The microbiome of the lichen *Lobaria pulmonaria* varies according to climate in Europe

**DOI:** 10.1101/2024.03.06.583579

**Authors:** Anteneh Tamirat Bogale, Maria Braun, Jörg Bernhardt, Daniela Zühlke, Ulf Schiefelbein, Manuela Bog, Christoph Scheidegger, Veronika Zengerer, Dörte Becher, Martin Grube, Katharina Riedel, Mia M. Bengtsson

## Abstract

The *Lobaria pulmonaria* holobiont comprises algal, fungal, cyanobacterial, and bacterial components. We investigated *L. pulmonaria’s* bacterial microbiome in the adaptation of this ecologically sensitive lichen species to diverse climatic conditions. Our central hypothesis posited that microbiome composition and functionality aligns with continental-scale climatic parameters related to temperature and precipitation. We also tested the impact of short-term weather dynamics, sampling season, and algal/fungal genotypes on microbiome variation. Metaproteomics provided insights into compositional and functional changes within the microbiome. Climatic variables explained 41.64% of microbiome variation, surpassing the combined influence of local weather and sampling season at 31.63%. Notably, annual mean temperature and temperature seasonality emerged as significant climatic drivers. Microbiome composition correlated with algal, not fungal genotype, suggesting similar environmental recruitment for the algal partner and microbiome. Differential abundance analyses revealed distinct protein compositions in sub-atlantic lowland and alpine regions, indicating differential microbiome responses to contrasting environmental/climatic conditions. Proteins involved in oxidative and cellular stress were notably different. Our findings highlight microbiome plasticity in adapting to stable climates, with limited responsiveness to short-term fluctuations, offering new insights into climate adaptation in lichen symbiosis.

## INTRODUCTION

Phenotypic plasticity, the ability of organisms to exhibit different characteristics under varying environmental conditions, is a key mechanism for coping with environmental changes (Bonamour et al., 2019). This adaptability enhances survival and reproductive success, contributing to fitness (Williams et al., 2017; Xue & Leibler, 2018). Moreover, phenotypic plasticity has the potential to moderate the loss of global biodiversity by enabling organisms to colonize new environments and undergo geographical range shifts (Bonamour et al., 2019). In holobionts, a term encompassing hosts and their associated microbial communities (Simon et al., 2019; Zilber-Rosenberg & Rosenberg, 2008), phenotypic plasticity can be complemented and supported by diverse microbial communities. These communities contribute functions that facilitate local environmental adaptation, such as supplying nutrients, providing resistance against abiotic factors, producing growth hormones, and facilitating atmospheric nitrogen fixation (Grimm et al., 2021; Kolodny & Schulenburg, 2020; Voolstra & Ziegler, 2020). Furthermore, the function and abundance of these host-associated and beneficial microbial communities are partly shaped by the environment and are associated with the microbial community’s fitness in that particular environment, resulting in new adaptive physiological functions for the host (Kolodny & Schulenburg, 2020).

Lichens, originally thought to be simple symbiotic relationships between fungi and algae (and/or cyanobacteria), are now recognized as complex holobionts, housing a diverse microbiome comprising bacteria, yeasts, protists, and viruses (Aschenbrenner et al., 2016; Grimm et al., 2021; Hawksworth & Grube, 2020; Simon et al., 2019). This microbiome may be important for enabling lichens to thrive in diverse terrestrial environments and adapt to shifting climatic conditions, facilitating nutrient acquisition and enhancing stress tolerance (Aschenbrenner et al., 2016; Grimm et al., 2021; Grube et al., 2015). This adaptability is especially crucial since long-term climatic factors, such as annual precipitation and temperature, determine regional climate patterns (Giordani & Incerti, 2008; Nascimbene et al., 2016), which may limit the distribution of the lichen. Meanwhile, short-term weather fluctuations introduce dynamic environmental challenges (Di Nuzzo et al., 2022a), which continually test the holobiont’s adaptability to sudden changes in hydration and light. Further, genotypic variation in the fungal and algal partners may play a role in shaping the microbiome, thus influencing the ecological strategies and adaptability of individual holobionts (Devkota et al., 2019; Fontaine et al., 2013; Ruprecht et al., 2012). An integrated investigation into the lichen microbiome is imperative not only for deepening our comprehension of this intricate symbiosis but also for illuminating how lichen holobionts and their associated microbial communities respond to environmental variations.

Metaproteomes, offering insights into the function, abundance dynamics, metabolism, and physiology of microbiomes, are valuable tools for understanding microbial communities (Kleiner, 2019; Salvato et al., 2021). Using metaproteomics to investigate microbiomes circumvents common culprits of DNA-based methods, as proteins reflect the current physiological state of cells, and not just their genetic potential. In this context, metaproteomics have led to breakthroughs in the study of symbiosis of selected holobiont systems, such as deep sea tube worms (Hinzke et al., 2021), lucinid clams (König et al., 2016) and the rice rhizosphere (Knief et al., 2011). First metaproteomics investigations into the microbiome of the model lichen species, *Lobaria pulmonaria* (L.) Hoffm. revealed possible roles of the microbiome in supplying nutrients, providing resistance against both biotic and abiotic stress, aiding in photosynthesis, supporting the growth of the fungal and algal partners, and possessing capabilities for detoxification and thallus degradation (Grube et al. 2015, Eymann et al. 2017). This large foliose lichen has been found to have associations with bacteria of high diversity (Aschenbrenner et al., 2016; Eymann et al., 2017; Grimm et al., 2021; Grube et al., 2015; Schneider et al., 2011). Compared to many other lichens, *L. pulmonaria* is less tolerant of desiccation and highly sensitive to air pollution (Grimm et al., 2021). However, the species has a worldwide distribution in diverse climatic regions, such as boreal, alpine, temperate, Mediterranean, and desert climates, which might be indicative of an association with a variable microbiome, lending plasticity to the holobiont (Aschenbrenner et al., 2016; Grimm et al., 2021; Grube et al., 2015). This unique combination of sensitivity to environmental stress on the one hand, and wide geographical distribution on the other hand, makes *L. pulmonaria* an interesting model holobiont for studying climate-related microbiome variation. However, there is currently little known about how the *L. pulmonaria* microbiome (or the microbiome of any other lichen species) varies according to climatic gradients.

In this work, we address the role of climatic variables on a continental scale in shaping microbiome composition and function in *L. pulmonaria*. We hypothesized that 1) climatic variables related to temperature and precipitation are the main drivers of bacterial microbiome metaproteome composition. In addition, we expected that 2) functional protein composition in the microbiomes reflects an adaptation of the lichen holobiont to climatic conditions. As alternative explanatory factors for microbiome variability, we included proxies of short-term weather and seasonality as well as genotypic information from the fungal and algal symbiotic partners. To address this, 42 specimens of the lung lichen *L. pulmonaria*, were collected from 14 different sampling locations in 5 countries in Europe and were analyzed using metaproteomics.

## EXPERIMENTAL PROCEDURES

### Sample collection and acquisition of climatic and local weather data

In September 2015, between May and July in 2017, and from October to November in 2018, fresh thalli of *Lobaria pulmonaria* (L.) Hoffm. were collected from trees in 14 different European sites. These sites included Koralpe and Tamischbachgraben in Austria (metaproteomics only), Lihme, Rebild, Rold Skov, and Viborg in Denmark, Johannishus, Kullen, Söderåsen, Stensnäs, and Vånga (metaproteomics only) in Sweden, Darß and Rhön in Germany, and Stara Rzeka in Poland. The samples were preserved in a cool box with thermal packs during transport and then stored at -70°C until protein extraction.

To assess the influence of climatic and local weather conditions on the *L. pulmonaria* microbiome, we gathered long-term climate data from the WorldClim database. From the original 19 bioclimatic layers, six with the lowest Pearson correlation coefficient (r ≤ 0·7) (AMT: Annual Mean Temperature, MDR: Mean Diurnal Range (Temperature), TS: Temperature Seasonality, AP: Annual Precipitation, PS: Precipitation Seasonality, and PDQ: Precipitation of Driest Quarter) were selected (Dormann et al., 2013).

In addition, short-term weather data, encompassing parameters like relative humidity, temperature, and precipitation cover for the two-week period leading up to each sampling event at each site (with a horizontal resolution of ½° x ⅝° latitude/longitude grid), was acquired from the NASA Prediction of Worldwide Energy Resources (NASA/POWER: https://power.larc.nasa.gov/data-access-viewer/) database. Since samples were taken at different times of the year, all sampling dates were converted into Julian dates, which was used as a proxy for sampling season.

### Protein extraction, mass spectrometry, and database search

Before protein extraction, we carefully removed visible external organisms like leaves, moss, and bark using sterile tweezers in order to avoid non-lichen contaminants. Subsequently, 300 mg of lichen thalli was rapidly shock-frozen with liquid nitrogen. Protein extraction was carried out as described previously (Eymann et al., 2017; Wang et al., 2006).

Aliquots of the extracted proteins were separated on an SDS polyacrylamide gel and stained overnight with Colloidal Coomassie Brilliant Blue. Subsequently, 15 protein bands were excised from the gel and digested with trypsin to generate peptides. These peptides were then extracted, and the resulting peptide mixtures from three technical replicates of each sample were analyzed by LC-MS/MS. The raw files were converted to MGF files using Proteome Discoverer and searched with the Mascot search engine. Scaffold software was utilized to validate MS/MS-based peptide and protein identifications. In the initial step, mass spectra from all samples were searched against the custom database. It was created combining the bacterial microbiome part of the database used in Eymann et al., (2017), a partly translated metagenome of *L. pulmonaria*, as well as publicly available genomic data from the taxa in Supplementary Table 1, downloaded from GenBank. The mass spectrometry proteomics data have been deposited to the ProteomeXchange Consortium via the PRIDE (Perez-Riverol et al., 2022) partner repository with the dataset identifier PXD049079.

### Functional and taxonomic annotation

Functional and taxonomic classifications of the proteins were conducted using the metaproteome analysis pipeline Prophane 2.0 (http://www.prophane.de, (Schiebenhoefer et al., 2020). To streamline the analysis, proteins were grouped based on shared peptide spectrum matches (PSMs). Within each protein cluster, a master protein was designated, selected based on comprehensive PSM coverage and probability scores. Quantification of protein abundance relied on the normalized spectral abundance factor (NSAF) values (Eymann et al., 2017). This approach allowed for a detailed assessment of protein functions and taxonomic assignments while reducing the complexity of the dataset, enhancing the precision of the analysis, and enabling meaningful comparisons among the protein groups.

### Protein filtering and data imputation

In proteomics, missing values (MVs), whether missing at random (MAR) or missing not at random (MNAR), can have a significant impact on the completeness and quality of the data, with MNAR being more commonly encountered and typically associated with abundance-dependent patterns. (Jin et al., 2021; Wei et al., 2018).

To address this issue, we evaluated the accuracy of various imputation methods to select the most suitable approach for our dataset. We started by filtering a total of 1785 proteins that were present in at least 2 out of 3 biological replicates across all sampling sites. A complete dataset (n=335) with no missing values was created for simulation. We introduced missing values (MNAR) into this complete dataset at three different rates (10%, 20%, 30%). These missing values were imputed using mean, median, random forest (RF), singular value defunction (SVD), and k-nearest neighbors (kNN) methods, implemented with the *impute_wrapper* functions in R. The performance of these imputation methods was evaluated based on normalized root mean squared error (NRMSE) and NRMSE-based sum of ranks (Jin et al., 2021; Wei et al., 2018). Among these methods, random forest (RF) exhibited the best performance, with the lowest NRMSE and NRMSE-based sum of ranks (Supplementary Figure 1). This method was used to impute the entire dataset, and a subset of the data containing only bacterial proteins (n=217) was further analyzed.

### Differential expression analysis

To unravel the influence of key climatic gradients on microbiome function and function, we meticulously classified the sampling sites into four climatic clusters, each defined by its unique climatic characteristics. Within these distinct climatic categories, we conducted a differential protein abundance analysis, focusing on the two regions exhibiting the most pronounced climatic disparities (Fig. 1B): the Alpine (ALP) and Sub Atlantic Lowland (SAL) clusters.

**Fig. 1:**
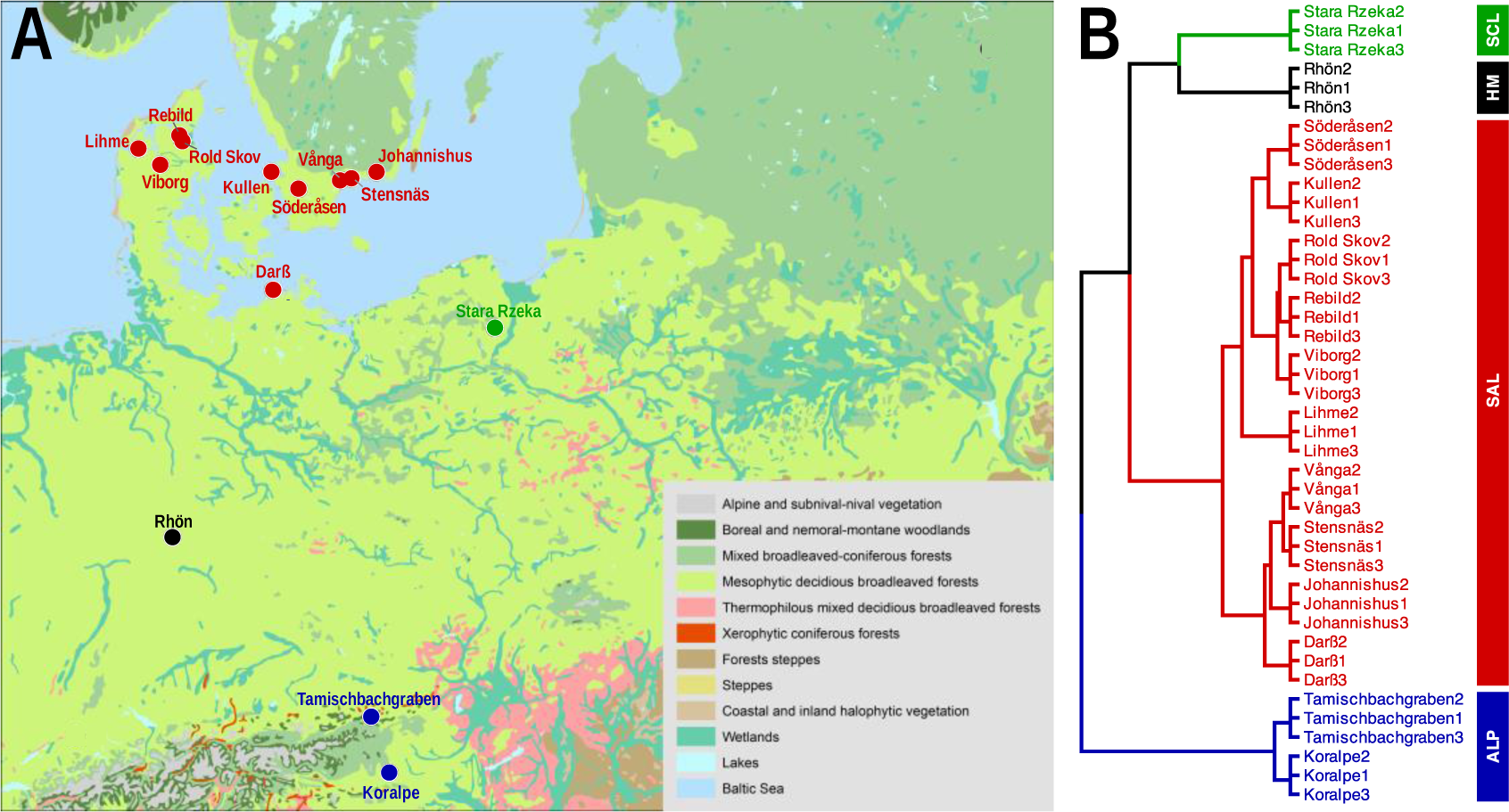
A map (**A**) showing the sampling sites across Europe: Koralpe and Tamischbachgraben (Austria); Lihme, Rebild, Roldskov, and Viborg (Denmark); Johannishus, Kullen, Söderåsen, Stensnäs, and Vånga (Sweden); Darß and Rhön (Germany); and Stara Rzeka (Poland), and a dendrogram (**B**) showing clusters of sampling sites based on climatic variables: Alpine (ALP), Sub Atlantic Lowland (SAL), Hercynian Montane (HM), and Sub Continental Lowland (SCL).

### Genotyping of fungal and algal partners

In our genotyping study, we systematically examined 46 samples from various climatic regions (not including the ALP region). This comprehensive analysis included the successful amplification of genetic markers using eight fungus-specific SSR loci and 14 algae-specific SSR loci (Supplementary Table 2), offering insights into the genetic diversity of *L. pulmonaria* lichens (Dal Grande et al., 2010; Walser et al., 2003). Our investigation into the assessment of genetic diversity within this symbiotic partnership commenced with the extraction of high-quality genetic material (Dal Grande et al., 2010; Scheidegger et al., 2012). Subsequently, we performed SSR analysis on both the fungal and algal components (Dal Grande et al., 2010; Widmer et al., 2012), targeting specific genetic loci characterized by repetitive DNA sequences to evaluate genetic variations. The result of this analysis consisted of scored fragments, which acted as markers for allelic distinctions. These scored fragments were then subjected to K-means clustering (Kamvar et al., 2014), allowing us to check the correspondence of the cluster results from the metaproteomic analysis with the genetic structure (Dal Grande et al., 2010; Walser et al., 2003).

### Statistical analysis

All statistical analyses were carried out in R (R Core Team, 2022). In order to describe the different climatic regimes of the sampling locations, we categorized our sites into four climatic clusters using a hybrid clustering approach combining hierarchical- and k-means (factoextra package) clustering based on the selected six non-redundant climatic variables (Kassambara & Mundt, 2020). These clusters, which also matched climatic classifications from the International Federation for Documentation (FID, 1971), were visualized using the ggpubr package’s *fviz_dend* function (Kassambara, 2023).

We used non-metric multidimensional scaling (nMDS) with a Bray–Curtis distance matrix to visualize site clustering based on metaproteome composition. The *envfit* function of vegan package was used to visualize the influence of climatic- and short-term weather variables on metaproteome composition. We then quantitatively assessed the impact of climatic and local weather variables on the microbiome through a permutation multivariate analysis of variance (PERMANOVA) with the *adonis2* function (Oksanen et al. 2022).

In order to identify differentially expressed proteins (DEPs) between the contrasting SAL and ALP regions, we used the Limma package (Ritchie et al., 2015). DEPs were considered significant if their adjusted p-value was <0.05, controlled for the false discovery rate (FDR) using the Benjamini-Hochberg procedure.

For assessing genetic diversity in *L. pulmonaria*, we imported scored fragment lengths into R and utilized the poppr v2.9.3 package (Kamvar et al., 2014) to categorize samples into three clusters through K-means clustering with the *find.clusters* function. We then performed a Mantel test (mantel function in the vegan package), employing Bray-Curtis dissimilarity and Spearman correlation) in order to assess the correlation of algal, fungal, and combined genotype profiles with microbiome composition (metaproteomic data). Bray-Curtis dissimilarity was calculated (vegdist, vegan package) based on the abundance of fungal and algal multilocus haplotypes (MLHf and MLHa) and their corresponding genetic clusters (GCf and GCa), as well as the abundance of metaproteomes across the sampling sites. Only sites represented in both datasets were included (i.e., excluding the ALP sites).

## RESULTS

### The *L. pulmonaria* microbiome: Functional and taxonomic insights

In our investigation of the microbiome using the Prophane metaproteomic pipeline, we explored its functional and taxonomic characteristics, revealing its versatility across various sampling locations. We identified a total of 217 unique proteins, which we classified into four distinct eggNOG functional categories (Fig. 2B). An average of 38.93% of the proteins were associated with “Metabolism,” with a particularly related to “Carbohydrate Transport and Metabolism “ and “Energy Production and Conversion “. Another 29.67% of the proteins played roles in “Information Storage and Processing,” primarily related to “Translation, Ribosomal Structure and Biogenesis.” Furthermore, 25.54% of the proteins were linked to “Cellular Processes and Signaling,” with a significant amount related to Post-Translational Modification, Protein Turnover, and Chaperones,. An intriguing category, denoted as “Poorly Categorized,” represented 5.86% of the proteins, proteins with insufficiently defined functions and characteristics.

**Fig. 2:**
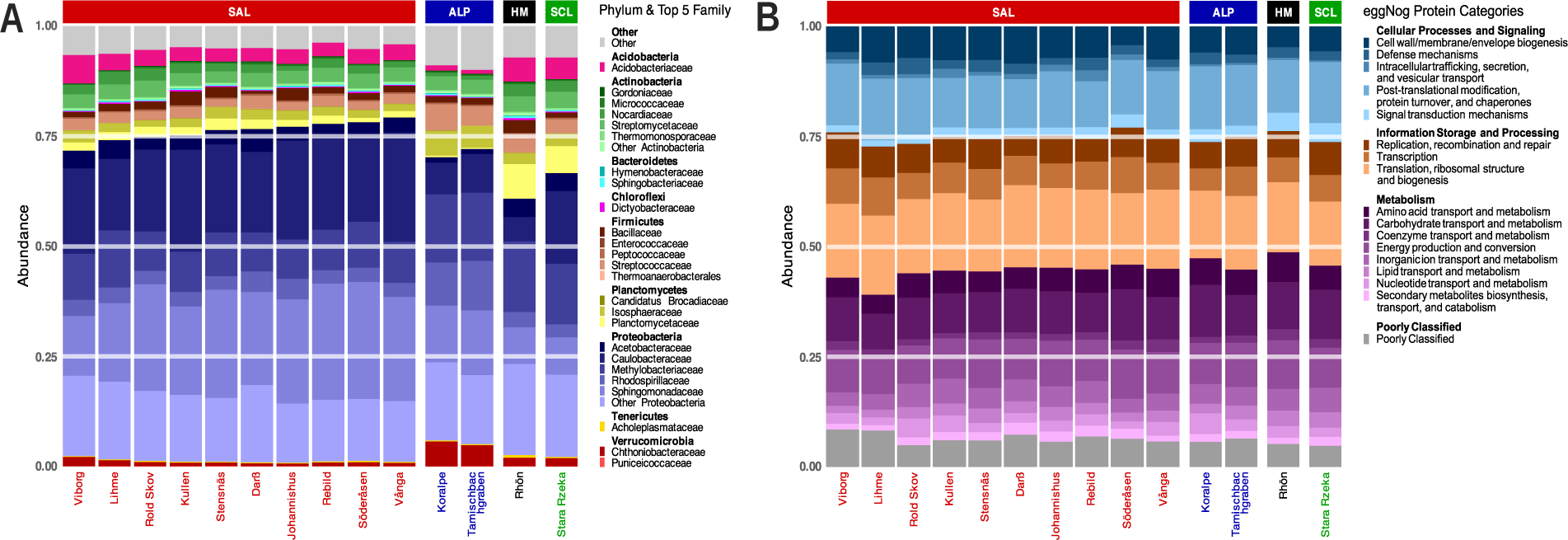
Taxonomic composition across sampling sites and climatic regions. When families are more than five in phylums, “Other + PHYLUM N ME” was shown, and stacked bars were sorted according to the Proteobacterial abundance for each sampling site (**A**) Functional Composition across sampling sites and climatic regions shown as eggNog main and sub role protein functional categories **(B)**.

Furthermore, our taxonomic analysis at the family level (Fig. 2A) revealed the presence of dominant microbial taxa consistently observed across the sampling sites. Notable families such as *Acidobacteriaceae, Streptomycetaceae, Caulobacteraceae, Methylobacteriaceae*, and *Sphingomonadaceae* were identified across these locations. At the phylum level, Proteobacteria emerged as the most dominant taxon. The relative abundance of Proteobacteria was shown to fluctuate among different families depending on sampling site. For instance, the Caulobacteraceae family ranged from 2.3% to 10.3% in the Rhön and Vånga sites, Methylobacteriaceae exhibited percentages between 5.6% and 10.4% in Darß and Rhön, and Sphingomonadaceae displayed percentages varying from 3.1% to 10.0% in the Rhön and Söderåsen sampling locations.

### Climate and weather impact on the *L. pulmonaria* microbiome metaproteome

The metaproteome composition of the *L. pulmonaria* microbiome was visualized in an NMDS plot highlighting the the variability between the sampling sites across Europe (Fig. 3A). The sites roughly clustered according to the climatic regions defined based on the climatic variables (Fig. 1B). Fitted vectors (arrows) representing both climatic data and short-term weather data, including sampling season in Julian date indicated that especially temperature seasonality (TS) and relative humidity 2 weeks prior to sampling (H2W) correlated with sample variability. Subsequently, a PERMANOVA analysis was conducted (Supplementary Table 4 and Supplementary Table 5), revealing the statistical significance of individual climatic and short-term weather variables. Notable climatic variables, such as Temperature Seasonality (TS) and Annual Mean Temperature (AMT), demonstrated significant influence cummulatively explaining 25.46% variation, with respective p-values of 0.001. The two most important short-term weather variables 2 weeks prior to sampling were relative humidity (H2W) and temperature (T2W) cumulatively explaining 20.12% (p-values <0.002). All together, the cumulative proportions explained by both data sets shed light on their relative importance. Climatic data collectively accounted for a substantial 41.64% of the variation in the microbiome metaproteome, highlighting its primary role in shaping the microbial function within *L. pulmonaria*. In comparison, short-term weather data, despite its significance, explained 31.63% of the variation.

**Fig. 3:**
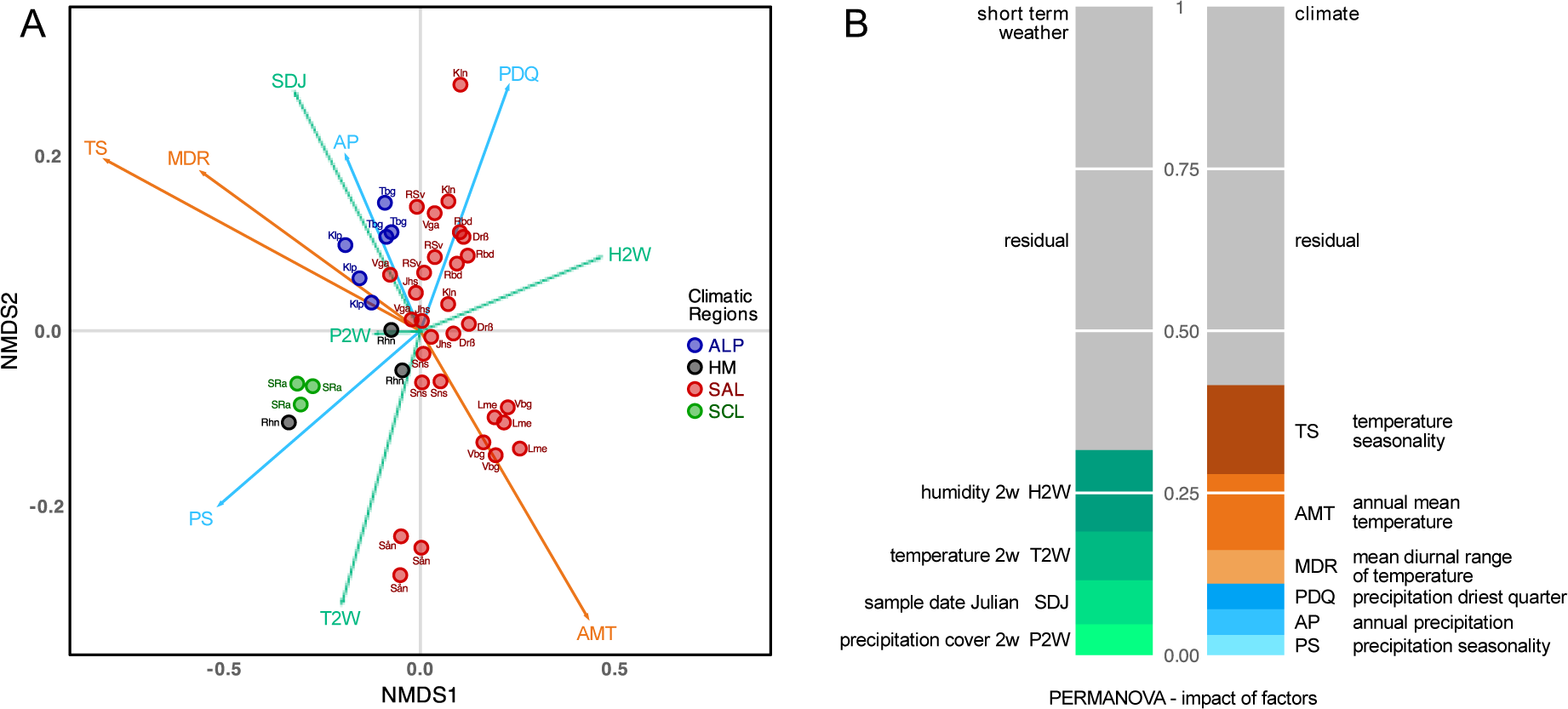
The influence of long-term climatic variables and short-term weather and seasonality on microbiome function across the sampling sites (**A**) (Rhn: Rhön, SRa: Stara Rzeka, Drß: Darß, Jhs: Johannishus, Kln: Kullen, Lme: Lihme, Rbd: Rebild, RSv: Rold Skov, Sån: Söderåsen, Sns: Stensnäs, Vga: Vånga, Vbg: Viborg, Klp: Koralpe, Tbg: Tamischbachgraben), and PERMANOVA analysis (**B**) on short-term weather (H2W, T2W, and P2W representing humidity, temperature and precipitation cover two weeks prior to sampling respectively, and SDJ: sample data in Julian) and climatic factors (TS, AMT, MDR, PDQ, AP and PS).

**Fig. 4:**
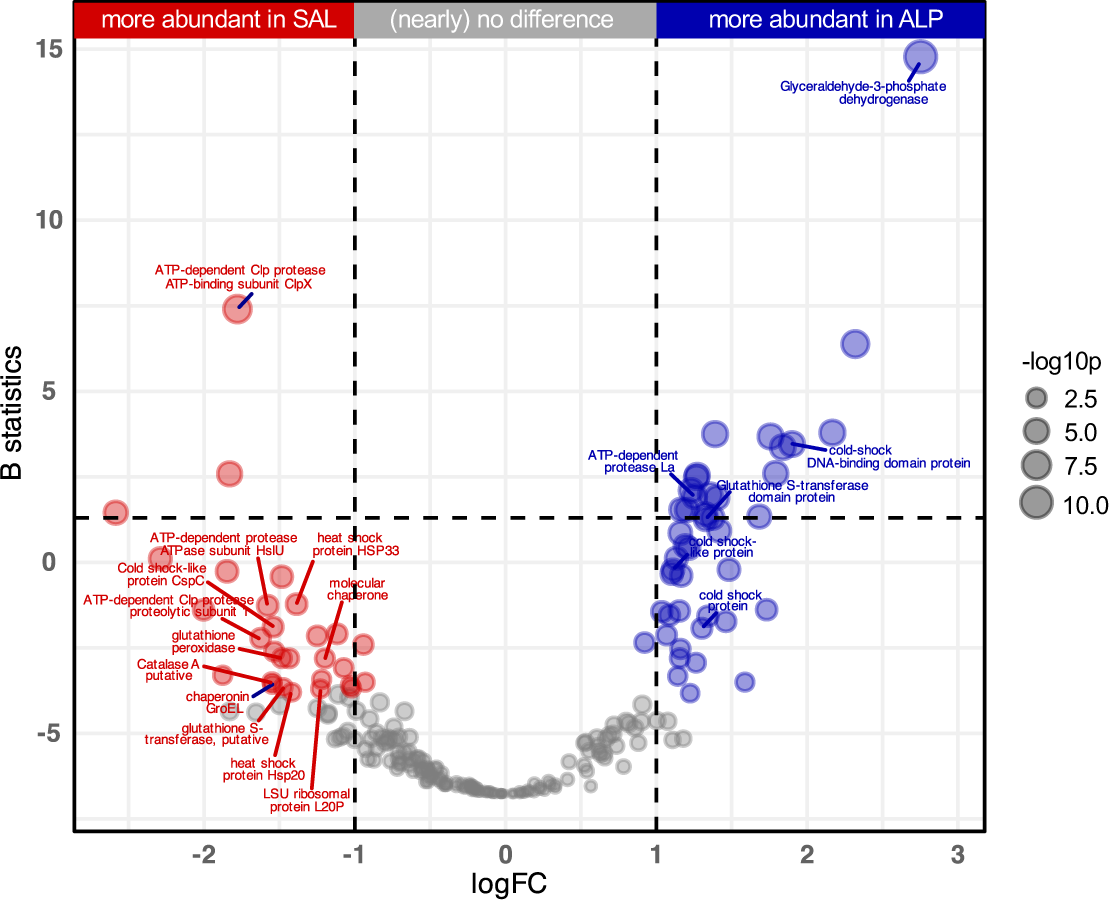
Differentially expressed proteins (DEPs) between SAL and ALP regions. Points before and after the line breaks represent increased abundance of proteins in ALP and SAL respectively. Y-axis represents B statistics (log-posterior odds of differential expression) and X-axis represents logFC (log2 of fold changes). Point diameter (-log10p) is proportional to -1*log10 (adjusted p value).

### Differential expression analysis

We performed a differential expression analysis on the most contrasting of our climatic regions, Sub-Atlantic Lowland (SAL) and Alpine (ALP) regions. In a differential abundance analysis, we identified 73 proteins, which showed distinct abundance patterns in these regions (44, more abundant in ALP and 29, more abundant in SAL). The SAL region exhibited increased levels of antioxidant proteins, including catalase A and glutathione peroxidase, crucial for mitigating oxidative stress by reducing reactive oxygen species (ROS) (Heck et al., 2010; Pei et al., 2023). Additionally, SAL displayed enhanced production of proteins associated with protein folding, refolding, and quality control, such as heat shock proteins and various proteases, indicative of active involvement in maintaining cellular integrity and responding to stress signals (Illigmann et al., 2021; Welch, 1993; Xie et al., 2013). In contrast, ALP exhibited elevated expression of Cold shock-like proteins CspG and CspC, which aid in mRNA synthesis and adaptation to low temperatures, as well as thioredoxin-related proteins like TrxA and thioredoxin reductase, essential for combating oxidative stress (Cardoza & Singh, 2022; Mariela et al., 2007). Furthermore, trehalose-related proteins were more abundant in ALP, including trehalose synthase and probable maltokinase, which are linked to desiccation protection (Leprince & Buitink, 2015; Tapia et al., 2015).

### Genotyping findings: Exploring genetic variation

In our genotyping analysis, a total of 46 samples representing various climatic regions (not including the ALP region), were systematically examined. We successfully amplified genetic markers using 8 fungus-specific SSR loci and 14 algae-specific SSR loci (Dal Grande et al., 2010; Walser et al., 2003) (Supplementary Table 2), providing insights into the genetic diversity of *L. pulmonaria* lichens. Our analysis detected a total of 51 alleles for the fungal symbiont and 100 alleles for the algal symbiont. The number of alleles per locus exhibited notable variation, ranging from 2 to 16 for the fungal partner and from 2 to 20 for the algal partner (Supplementary Table 2). By combining these allelic data into multi-locus haplotypes (MLHs), we identified 19 unique MLHs for the fungal partner and 30 unique MLHs for the algal partner (Supplementary Table 3), underscoring the genetic complexity of these symbiotic associations. Furthermore, our analysis uncovered different MLHs within 5 out of 12 localities for the fungal partner and within 9 out of 12 localities for the algal partner, highlighting the pronounced genetic heterogeneity across the diverse geographic locations. Interestingly, various combinations of fungal and algal MLHs were observed within several localities, contributing to the overall genetic variability of the lichen. Samples assignable to different genetic clusters were distributed within the two climatic clusters of regions HM and SAL for both the fungal and algal partners. Additionally, samples from the Rhön and Darß localities exhibited dual membership in two of the genetic clusters. This reflects the complex phylogeographic colonization processes in *L. pulmonaria* which resulted in an intricate genetic geographic structure in Europe (Lerch et al., 2018; Nadyeina et al., 2014; Scheidegger et al., 2012; Widmer et al., 2012). The Mantel test results comparing algal, fungal and combined genetic profiles (clusters) in relation to microbiome composition (metaproteomic data) revealed fair correlations. The combined genotypic dataset showed a moderate correlation with the microbiome (Mantel R = 0.3509, p = 0.0272). In contrast, the fungal genotype profiles did not significantly correlate with the microbiome data (Mantel R = 0.02031, p = 0.4011). However, the algal genotype profiles displayed a moderate yet significant correlation with the microbiome (Mantel R = 0.3880, p = 0.0293).

## DISCUSSION

In this study, our primary objective was to gain insight into how the *L. pulmonaria* microbiome reacts to ecological factors, with a specific focus on climatic conditions, short-term weather dynamics and sampling season as well as genotypic diversity of fungal and algal symbiotic partners. We used metaproteomics to simultaneously address both taxonomic and functional variation of the microbiome. Our analysis uncovered the significant impact of long-term climate-related factors, particularly temperature-related variables, on the *L. pulmonaria* microbiome. However, short-term weather dynamics also explained a significant amount of variation. Furthermore, our exploration of functional gene expression revealed noteworthy differences in protein profiles between two contrasting climatic regions, SAL and ALP. Our investigation into genetic diversity unveiled an intricate genetic structure of *L. pulmonaria* fungal and algal partners, yet this diversity did not align with the metaproteome profiles of the microbiome. These findings may contribute to a deeper understanding of the adaptability and resilience of this lichen holobiont in the face of climate change.

### The role of climate in shaping *L. pulmonaria* microbiome

Our analysis of *L. pulmonaria*’s metaproteome structure suggests the dominant influence of long-term climatic variables as a driver of microbiome taxonomic composition and functionality. We hypothesized that factors related to temperature and precipitation would be the main drivers of the microbiome metaproteome composition. Our findings singled out particularly temperature-related factors, including temperature seasonality and annual mean temperature as important, jointly explaining 25.46% of the microbiome variation. Temperature Seasonality, responsible for 13.74% of the variation, is a measure of the annual range in temperature. A high seasonal temperature fluctuation may necessitate a high functional plasticity of the microbiome to sustain microbial metabolic activity within lichen thalli at various temperatures. Annual Mean Temperature, contributing 11.72% to the microbiome variation, regulates resource availability, primary productivity (Pierre et al., 2020), and microbial interactions (Apple et al., 2006; Knapp & Huang, 2022). The influence of these variables on lichen physiology, growth, and metabolism, including their effects on metabolic rates, water availability, and nutrient dynamics, aligns with already existing research (Finger-Higgens et al., 2022; Ladrón de Guevara et al., 2018; Mallen-Cooper et al., 2023; Nascimbene et al., 2016). Notably, while temperature-related variables emerge as key contributors, precipitation-related variables, such as precipitation seasonality and annual precipitation, play a comparatively minor role. This may indicate potential limitations in the representation of microclimatic conditions which may be more relevant for the individual lichen holobiont than large-scale precipitation and humidity regimes. Based on this, the need for a more comprehensive understanding of precipitation- and humidity-related factors and their impact on microbial communities in lichens is evident. High-resolution, localized data collection and monitoring are essential for gaining precise insights into microclimate.

### The role of short-term weather and seasonality for microbiome dynamics

A notable challenge in microbiome field studies is capturing the relevant time scale at which dynamics in composition and functional gene expression occur. Microorganisms have short generation times, and can thereby respond rapidly to changing environmental conditions, leading to compositional and functional variability on a short-term time scale (Di Nuzzo et al., 2022b; Gauslaa, 2014). We therefore suspected that the weather immediately before the time of sampling, as well as the season of sampling, could have a large impact on microbiome composition, potentially comparable to the long-term prevailing climate at the sampling site.

In order to capture potential short-term compositional and functional variability, studying the effects of weather data directly in connection with sampling is important. However, when direct measurements of weather-related parameters are not available or are scarce, satellite-based observations can be considered (Marzouk, 2021; Rodrigues & Braga, 2021). We used POWER (Prediction Of Worldwide Energy Resources) by NASA which is generally contiguous in time, containing a global grid with a resolution of 0.5 latitude by 0.5 longitude. Despite this limited resolution, our findings showed that, although explaining less variation (31.63%) compared to long-term climatic factors, short-term local weather also likely plays a role in shaping *L. pulmonaria* microbiome community composition. Interestingly, we observed that relative humidity during the last 2 weeks prior to sampling explained the most variation (12.60%) while temperature and precipitation cover appeared less important which is in contrast to the greater importance of temperature-related climatic factors. This may indicate that the microbiome responds quickly to changes in relative humidity, while temperature regimes put longer-term constraints on microbiome composition. Short-term dynamics in microbiome composition are relatively poorly understood even for relatively well-studied model organisms (Aptroot & van Herk, 2007; Gerber, 2014; Kuzyakov & Razavi, 2019; van Oppen & Blackall, 2019), and our findings further highlight the need for experimental approaches that capture these dynamics at a higher temporal resolution and with more accurate measurement of environmental parameters than we could achieve here. A further limitation is caused by the fact that climate and weather are not independent from each other. Despite short-term fluctuations, weather is dependent on climate, making it difficult to separate the impact of one or the other on microbiome dynamics.

### The role of genotypic variation in the fungal and algal symbiotic partners

We carried out genetic analyses on lichen material collected from a specific set of sampling sites, generating genetic profiles for the primary symbiotic partners of the lichen holobiont. The genotyping analysis, detailed in Supplementary Table 2, revealed significant genetic diversity. To be more specific, the fungal symbiont exhibited 51 alleles, forming 19 haplotypes, and the algal symbiont presented 100 alleles, resulting in 30 haplotypes. This genetic complexity was evident in 5 out of 12 fungal and 9 out of 12 algal localities.

Despite the genetic variability observed in both symbiotic partners of the holobiont, our microbiome (metaproteome) data only correlated moderately with the algal genotype composition, as indicated by our mantel test results. The correlation between algal genotype and microbiome composition is consistent with the recruitment of both the algal partner and the microbiome from the local environment following sexual reproduction and dispersal of the fungal partner. In *L. pulmonaria*, clonal reproduction, when vegetative propagules containing both fungal, algal and presumably microbiome components are dispersed over shorter distances is also important (Grimm et al. 2020). Additionally, the presence of recurring multilocus genotypes in *L. pulmonaria* influenced the spatial genetic structure solely within low-density populations (Walser et al., 2004). Altogether, this supports the idea of local adaptation of the holobiont via recruitment of the bacterial microbiome from the environment, consisting of bacterial taxa adapted to the prevailing climatic conditions at the site.

While our study highlights a correlation between microbiome composition and algal genotype, there is a limitation due to the exclusion of Alpine sampling sites in the genotyping analysis (Supplementary Table 3). Looking ahead, we propose a comprehensive examination covering all sampled sites to thoroughly evaluate the alignment between microbiome phenotypes and genotypes in *L. pulmonaria*. Such an inclusive investigation will not only confirm the consistency of observed correlations across diverse sites but will also unveil the relative contributions of clonal propagation and sexual recombination in shaping microbial communities.

### Molecular mechanisms of microbiome adaptation

We explored the functional gene expression in the microbiome of *L. pulmonaria*, unveiling its adaptation to the distinct climatic conditions of the Sub-Atlantic Lowland (SAL) and Alpine (ALP) regions. These regions, defined by pronounced differences in the two most important climatic variables identified in our study, average mean temperature (AMT) and temperature seasonality (TS), featured significant differences in the holobiont’s proteomes, further supporting the role of climate in shaping the microbiome. ALP was characterized by high TS, while SAL was characterixzed by lower TS and higher AMT (Supplementary Table 6). We hypothesized that functional protein composition in the microbiomes reflects an adaptation of the lichen holobiont to climatic conditions. Indeed, comparing SAL and ALP, we observed differential protein abundances indicative of adaptation to their specific environments. SAL, characterized by higher AMT and lower TS, exhibited an abundance of proteins addressing oxidative stress. Notably, catalase A and glutathione peroxidase were prominent, which are linked to decomposing hydrogen peroxide and eliminating reactive oxygen species (Heck et al., 2010; Pei et al., 2023). Additionally, ribosomal proteins were found to be increased in SAL, potentially indicating higher demands for protein synthesis, possibly related to a higher metabolic activity in this comparably warm region.

Conversely, ALP, with colder and drier conditions (Alps et al., 2001; Humphries, 2020) featured significantly more abundant proteins enhancing cold stress adaptation, such as Cold shock-like protein CspG and CspC, and thioredoxin-related proteins like TrxA and thioredoxin reductase, involved in response to oxidative stress (Cardoza & Singh, 2022; Mariela et al., 2007). Furthermore, proteins addressing heat and oxidative stress, including heat shock proteins, ATP-dependent Clp protease proteolytic subunit 1, and ATP-dependent protease ATPase subunit HslU, were more abundant in SAL (Illigmann et al., 2021; Welch, 1993; Xie et al., 2013). In ALP, trehalose-related proteins, trehalose synthase, and maltokinase, involved in adaptation to desiccation (Leprince & Buitink, 2015; Tapia et al., 2015) were more abundant.

It is plausible that the apparent proteomic responses of the microbiome to the prevailing climate contribute adaptability of the whole lichen holobiont through interactions between bacterial microbiome and the fungal and algal partners (Grimm et al., 2021). For example, trehalose-producing microorganisms have been suggested to also lend desiccation tolerance to plant hosts (Vílchez et al., 2016). Similarly, certain bacteria have been shown to improve the antioxidant status of plants grown under drought stress (Chiappero et al., 2019; De Vries et al., 2020). However, the interaction mechanisms by which microbiome bacteria may transfer e.g., secondary metabolites to the other symbiosis partners are difficult to disentangle in an observational study such as ours. A targeted experimental approach by which the holobiont is subjected to defined stressors, while monitoring the gene expression of the microbiome, fungal and algal partners would be ideal to tackle these central questions.

## CONCLUSIONS

Our research on the microbiome of *L. pulmonaria* (L.) Hoffm has unveiled the pivotal role of climate, particularly annual mean temperature (AMT) and temperature seasonality (TS), in shaping the microbiome of *L. pulmonaria*. These climatic factors predominantly explain microbiome variation, underlining the profound influence of long-term environmental conditions. While short-term weather and seasonality appeared to play a secondary role, they likely contribute to the microbiome’s function, and highlight the need to take into account dynamics on different temporal scales in microbiome studies.

Moreover, our study highlights the microbiome’s remarkable adaptability to diverse climates and environments, as evidenced by differentially abundant proteins and their functions, including oxidative stress mitigation, ROS scavenging, and desiccation tolerance. We found that algal genotype correlates with microbiome composition, consistent with a similar mode of local recruitment for the algal partner and the microbiome. This research underscores the broader significance of the microbiome in aiding holobiont adaptation, although the specific mechanisms by which the microbiome may aid the entire holobiont in climate adaptation remain elusive.

Moving forward, we recommend further research employing experimental approaches with higher temporal resolution and more precise measurements of environmental parameters to enhance the accuracy and granularity of data. Such approaches, especially combined with controlled manipulation of environmental stressors, will yield deeper insights into *L. pulmonaria*’s microbiome dynamics and its response to its surroundings. In addition, studying the microbiome of *L. pulmonaria* over wider climatic gradients and in other geographic regions, profiting from the broad global distribution of this lichen species, may confirm or challenge our conclusions on the role of climate for shaping the microbiome structure and function.

## ACKNOWLEDGEMENTS

We acknowledge the financial support for this research provided by the Deutsche Forschungsgemeinschaft (DFG) through the Research Training Group ‘Biological RESPONSEs to Novel and Changing Environments’ (GRK 2010), subproject A4n “The lichen microbiome and its role in adaptation to climate change associated factors”. We also extend our appreciation to all members of the GRK 2010 consortium for offering valuable feedback that significantly enhanced the quality of the data analysis. Special thanks are given to Prof. Dr. Jürgen Kreyling for his support. Finally, we express our thanks to Dr. Dirk Albrecht for his assistance in bioinformatics infrastructure.

## DATA AVAILABILITY

Data generated or analyzed during this study are available from the corresponding author upon reasonable request.

