## Supplemental Files for "The microbiome of the lichen *Lobaria pulmonaria* varies according to climate in Europe"

**LIST OF SUPPLEMENTARY TABLES AND FIGURES**

Supplementary Table 1: Families of NCBI taxonomy database included in the custom database used in this study.

| | | |
|---|---|---|
| Acetobacteraceae | Fimbriimonadaceae | Nocardiopsaceae |
| Acholeplasmataceae | Fonticulaceae | Nosematidae |
| Albuginaceae | Frankiaceae | Nostocaceae |
| Alphabaculovirus | Geminigeraceae | Oleaceae |
| Anguinidae | Gloeobacteraceae | Oxytrichidae |
| Archangiaceae | Glomeraceae | Papillomaviridae |
| Arteriviridae | Gordoniaceae | Patulibacteraceae |
| Bacteriovoracaceae | Hoplolaimidae | Peltigerales |
| Baculoviridae | Iridoviridae | Peptococcaceae |
| Basidiomycota | Isosphaeraceae | Phycisphaeraceae |
| Betaflexiviridae | Kineosporiaceae | Physaraceae |
| Brevibacteriaceae | Kofleriaceae | Planctomycetaceae |
| Bromoviridae | Ktedonobacteraceae | Polyangiaceae |
| Bryophyta | Lecanoromycetes | Prochloraceae |
| Candidatus Brocadiaceae | Lecanoromycetidae | Prolixibacteraceae |
| Cellulomonadaceae | Legionellaceae | Propionibacteriaceae |
| Chromobacteriaceae | Leuconostocaceae | Reoviridae |
| Competibacteraceae | Lichtheimiaceae | Retroviridae |
| Cryptosporangiaceae | Lobaria | Rhabdoviridae |
| Culicosporidae | Lobariaceae | Rhizopodaceae |
| Cystobacter | Malvaceae | Rhodospirillaceae |
| Cystobacteraceae | Marchantiophyta | Rubrobacteraceae |
| Dehalococcoidaceae | Methanomicrobiaceae | Salicaceae |
| Dermabacteraceae | Methylophilaceae | Sapindaceae |
| Desulfarculaceae | Micrococcaceae | Scytonemataceae |
| Dictyochloropsis | Moraxellaceae | Sinobacteraceae |
| DictyochloropsisReticulata | Mortierellaceae | Spiroplasmataceae |
| Eggerthellaceae | Mucoraceae | Staphylococcaceae |
| Elusimicrobiaceae | Myxococcaceae | Tardigrada |
| Entamoebidae | Nannocystis | Thermoleophilia |
| Enterococcaceae | Nitrosopumilaceae | Thermomonosporaceae |
| Erwiniaceae | Nitrososphaeraceae | Trebouxiaceae |
| Fagaceae | Nocardiaceae | Vavraia |

Supplementary Figure 1: Evaluation of Missing value imputation: Normalized root mean squared error (NRMSE), and NRMSE-based sum of ranks

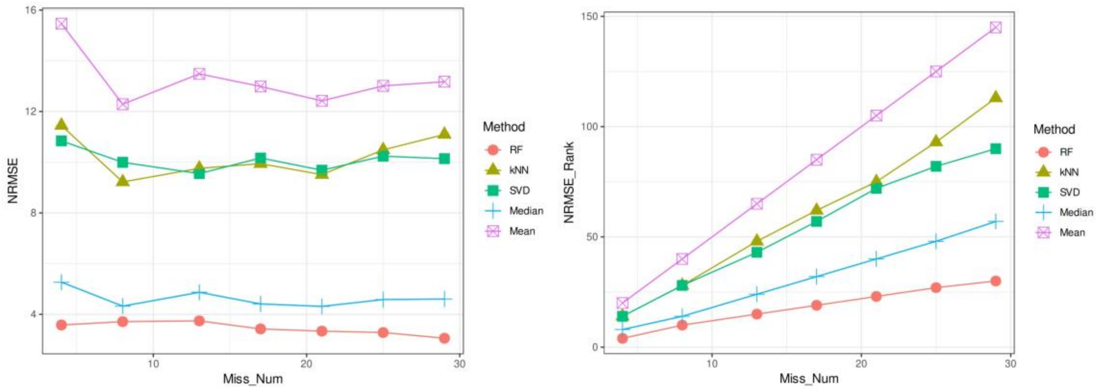

Supplementary Table 2: Symbiont-specific SSR markers. Number of alleles, allele size range and percent of missing data detected for the 46 investigated samples are given.

| | Number of alleles | Allele size range (bp) | % missing |
|---|---|---|---|
| Fungus-specific SSR markers | | | |
| LPu03 | 4 | 185-191 | 2.2 |
| LPu09 | 4 | 173-281 | 6.5 |
| LPu15 | 8 | 157-199 | 0.0 |
| LPu23 | 3 | 295-312 | 0.0 |
| LPu24 | 2 | 229-239 | 0.0 |
| LPu25_12 | 16 | 196-292 | 0.0 |
| LPu28 | 11 | 270-326 | 0.0 |
| MS4 | 3 | 184-238 | 0.0 |
| Algae-specific SSR markers | | | |
| 6816 | 9 | 160-231 | 0.0 |
| 6819 | 11 | 136-173 | 4.3 |
| 6820 | 7 | 197-214 | 0.0 |
| 6825_2 | 3 | 133-153 | 0.0 |
| 6528 | 4 | 102-108 | 0.0 |
| 6861_3 | 3 | 120-127 | 0.0 |
| 6863 | 7 | 147-198 | 0.0 |
| 7000_2 | 5 | 155-164 | 0.0 |
| 7007_2 | 5 | 171-179 | 0.0 |
| LPu16 | 9 | 196-212 | 0.0 |
| LPu19 | 2 | 441-444 | 30.4 |
| LPu20 | 12 | 190-254 | 0.0 |
| LPu26 | 20 | 354-513 | 23.9 |
| LPu27 | 3 | 194-198 | 0.0 |

Supplementary Table 3: Results of the SSR analysis for both symbiotic partners of *Lobaria pulmonaria* for the 12 investigated localities. Given are the unique multilocus haplotypes for the fungal and algal partner (uMLHf and uMLHa), the resulting genetic clusters from the cluster analysis for the two partners (GCf and GCa) and the corresponding climate region.

| Location | uMLHf | uMLHa | GCf | GCa | Climate Region* |
|---|---|---|---|---|---|
| Stara Rzeka1 | f1 | a1 | fA | aA | SCL |
| Stara Rzeka2 | f1 | a1 | fA | aA | SCL |
| Stara Rzeka3 | NA | NA | NA | NA | SCL |
| Rhön1 | f2 | a2 | fB | aA | HM |
| Rhön2 | f3 | a3 | fB | aB | HM |
| Rhön3 | f4 | a4 | fA | aB | HM |
| Darß | f5 | a5 | fB | aA | SAL |
| Darß | f5 | a5 | fB | aA | SAL |
| Darß | f5 | a5 | fB | aA | SAL |
| Darß | f5 | a5 | fB | aA | SAL |
| Darß | f6 | a6 | fC | aA | SAL |
| Darß | f6 | a6 | fC | aA | SAL |
| Darß | f6 | a7 | fC | aA | SAL |
| Darß | f7 | a8 | fC | aC | SAL |
| Darß | f7 | a8 | fC | aC | SAL |
| Darß | f7 | a9 | fC | aC | SAL |
| Lihme1 | f8 | a10 | fC | aC | SAL |
| Lihme2 | f9 | a10 | fC | aC | SAL |
| Lihme3 | f9 | a11 | fC | aC | SAL |
| Rebild1 | f10 | a12 | fB | aC | SAL |
| Rebild2 | f10 | a12 | fB | aC | SAL |
| Rebild3 | f10 | a13 | fB | aC | SAL |
| Rold Skov1 | f11 | a14 | fA | aC | SAL |
| Rold Skov2 | f11 | a14 | fA | aC | SAL |
| Rold Skov3 | f11 | a14 | fA | aC | SAL |
| Viborg1 | f12 | a15 | fC | aC | SAL |
| Viborg2 | f12 | a16 | fC | aC | SAL |
| Viborg3 | f12 | a15 | fC | aC | SAL |
| Johannishus1 | f13 | a17 | fA | aA | SAL |
| Johannishus2 | f14 | a18 | fA | aA | SAL |
| Johannishus3 | f14 | a17 | fA | aA | SAL |
| Kullen1 | f15 | a19 | fC | aC | SAL |
| Kullen2 | f16 | a20 | fC | aC | SAL |
| Kullen3 | f15 | a21 | fC | aC | SAL |
| Söderåsen | f17 | a22 | fC | aB | SAL |
| Söderåsen | f17 | a23 | fC | aB | SAL |
| Söderåsen | f17 | a24 | fC | aB | SAL |
| Söderåsen | f17 | a25 | fC | aB | SAL |
| Söderåsen | f17 | a26 | fC | aB | SAL |
| Stensnäs | f18 | a27 | fA | aC | SAL |

| | | | | | |
|---|---|---|---|---|---|
| Stensnäs | f18 | a28 | fA | aC | SAL |
| Stensnäs | f18 | a28 | fA | aC | SAL |
| Stensnäs | f18 | a29 | fA | aC | SAL |
| Stensnäs | f18 | a28 | fA | aC | SAL |
| Vånga1 | f19 | a30 | fB | aA | SAL |
| Vånga2 | f19 | a30 | fB | aA | SAL |
| Vånga3 | f19 | a30 | fB | aA | SAL |

HM – Hercynian Montane, SAL – Sub Atlantic Lowland, SCL – Sub Continental Lowland.

Supplementary Table 4: The microbiome variation explained by climatic variables according to PERMANOVA

| Bioclimate variables | Df | Sum Sq | $R^2$ | F | Pr(>F) |
|---|---|---|---|---|---|
| Temperature Seasonality (TS) | 1 | 0.43287 | 0.13743 | 8.2411 | 0.001 *** |
| Annual Mean Temperature (AMT) | 1 | 0.36912 | 0.11719 | 7.0274 | 0.001 *** |
| Mean Diurnal Range (Temperature) (MDR) | 1 | 0.16346 | 0.05189 | 3.1119 | 0.014 * |
| Precipitation of Driest Quarter (PDQ) | 1 | 0.12456 | 0.03954 | 2.3713 | 0.025 * |
| Annual Precipitation (AP) | 1 | 0.12443 | 0.03950 | 2.3689 | 0.038 * |
| Precipitation Seasonality (PS) | 1 | 0.09700 | 0.03079 | 1.8466 | 0.074 |
| Residual | 35 | 1.83840 | 0.58365 | | |
| Total | 41 | 3.14983 | 1.00000 | | |

Supplementary Table 5: The microbiome variation explained by variables relating to weather and season at the time of sampling according to PERMANOVA

| Local weather variables | Df | Sum Sq | $R^2$ | F | Pr(>F) |
|---|---|---|---|---|---|
| Relative Humidity (H2W) | 1 | 0.39663 | 0.12592 | 6.8140 | 0.001 *** |
| Temperature (T2W) | 1 | 0.23716 | 0.07529 | 4.0743 | 0.002 ** |
| Sampling Season (Julian Date, SSJ) | 1 | 0.21361 | 0.06782 | 3.6699 | 0.003** |
| Precipitation Cover (P2W) | 1 | 0.14874 | 0.04722 | 2.5554 | 0.023 * |
| Residual | 37 | 2.15369 | 0.68375 | | |
| Total | 41 | 3.14983 | 1.00000 | | |
